# Biglycan Promotes Glioma Cell Line Epithelial-Mesenchymal Transition and Temozolomide Resistance in Vitro

**DOI:** 10.1101/2024.03.19.581854

**Authors:** Xiangming Han, Yan Zheng

## Abstract

Gliomas are aggressive brain tumors associated with high mortality and treatment resistance. This study investigates the role of Biglycan (BGN) in glioma progression and its potential as a prognostic marker and therapeutic target. Analysis of data from The Cancer Genome Atlas (TCGA) reveals dysregulated BGN expression in low-grade gliomas (LGG) and glioblastoma (GBM) with positive correlation to tumor grade. High BGN expression is associated with poor prognosis. Functional analysis highlights BGN’s involvement in ECM regulation and EMT, driving glioma invasion and drug resistance. Silencing BGN reduces EMT-related gene expression and invasion capacity. Moreover, BGN expression is upregulated upon TMZ treatment, implicating its role in TMZ resistance. Targeting BGN may be a promising strategy to overcome glioma treatment resistance.

## Introduction

Glioma is a highly aggressive and fatal brain tumor with low 5-year survival rate[1]. The extracellular matrix (ECM), a complex network of proteins and carbohydrates surrounding cells, has been implicated in various aspects of glioma development and progression[2]. Specifically, the ECM has been associated with glioma cells undergoing epithelial-mesenchymal transition (EMT), a process that enhances tumor invasiveness and metastasis, as well as with the resistance of glioma cells to the standard chemotherapy drug, Temozolomide. These factors contribute to the poor prognosis and limited effectiveness of current therapeutic approaches [3,4]. One intriguing component of the ECM is biglycan (BGN), a secreted protein that interacts with various components of the ECM through its core protein or glycosaminoglycan chains[5,6]. Biglycan becomes sequestered within the ECM of most organs, suggesting its involvement in fundamental biological processes. Initially, biglycan was considered a static structural element of the ECM, providing physical support and stability. However, an increasing number of research articles have highlighted the potential role of biglycan in tumorigenesis, including its impact on cancer progression[7,8].Despite the growing interest in biglycan’s involvement in cancer, its precise role within the ECM, particularly in the context of glioma progression, remains largely unidentified. Investigating the influence of biglycan on glioma cells and the surrounding microenvironment could provide valuable insights into the mechanisms underlying glioma development, invasiveness, and therapeutic resistance. Understanding the interplay between biglycan and glioma may lead to the discovery of novel therapeutic targets or the development of innovative treatment strategies aimed at improving patient outcomes.

## Materials and methods

### 1. Patients and glioma cell lines

Clinical data from patients and RNA sequencing data of glioma samples were acquired from TCGA and CGGA databases. U251MG and HS683 cell lines were bought from American Type Culture Collection (ATCC, Gaithersburg, MD, USA), cultured in DMEM (Gibco, USA) containing 10% fetal bovine serum (FBS; Sijiqing, Hangzhou China) and 100 U/ml penicillin, 100 µM streptomycin (Gibco, USA) at 37 °C with 5% CO2.

### 2. Small interfering RNA (siRNA)

BGN non-targeting control siRNA(siNC) and siRNA from GenePharma Inc (Shanghai, China) were used to establish different BGN expression U251MG and HS683.

Transfection reagent Lipofectamine™ 3000 transfection reagent (Invitrogen, USA) was utilized to promote siRNA get into cells. The siRNA sequences targeting BGN are listed below:

siNC: forward, 5′-GCAUCAGCCUCUUCAACAATT-3′, reverse, 5′-UUGUUGAAGAG GCUGAUGCTT-3′, siBGN-a: forward 5′-GCCAUUCAUGAUGAACGAUTT-3′, revers e 5′-AUCGUUCAUCAUGAAUGGCTT-3′; siBGN-b: forward,5′-GCAUCAGCCUCUU CAACAATT-3′, reverse, 5′-UUGUUGAAGAGGCUGAUGCTT-3′; siBGN-c: forward 5 ′-GTC TAT CTG CAC TCC AAC AA-3′; reverse, 5′-TGG ATG GCC AGG CGG TCA GT-3′.

### 3. Western blot

Western blot analysis was performed as described previously[9], BGN (1:500, 16409-1-AP, Proteintech, USA), N-cadherin (1:200, sc-59987, Santa Cruz, USA), Vimentin (1:200, sc-373717, Santa Cruz, USA), β-actin (1: 5000, Cell Signalling Technology) were utilized in western blot to check invasive capacity of glioma cells. SRY-Box Transcription Factor 2(SOX2, Cell Signalling Technology) and β-catenin (Cell Signalling Technology) was used to test expression stemness marker of TMZ treated cells.

### 4. Transwell

As described previously[10], U251MG and HS683 (1 × 10^5^ cells) were seeded into the upper chamber (Corning, USA) precoated with Matrigel (Corning, USA). The lower chamber was filled with 500□μl of DMEM containing 10% FBS. Transwell chambers were placed in a 37□°C incubator with 5% CO2 for 24□hours. Subsequently, cells were fixed with 4% paraformaldehyde, stained with crystal violet (Sigma-Aldrich) and images were captured using Carl Zeiss Microscopy with both 1x and 10x objective magnification.

### 5. Colony formation assay

As described previously[11], cells were treated with 200 µM Temozolomide for 4 hours, 48 hours after siRNA silencing. Subsequently, the cells were seeded in 10 cm dishes (Corning, USA) at a concentration of 800 cells per dish in fresh complete media. After 10 days, the cells were fixed using 4% paraformaldehyde and stained with crystal violet. The number of colonies containing at least 50 cells was quantified to assess the effect of siRNA silencing and TMZ treatment on colony formation.

### 6. Statistical analysis

GraphPad Prism version 9.0 software was used for graph preparation and data analysis. Densitometric analysis was performed by ImageJ software. The significance of differences was analyzed using a t-test or one-way ANOVA and P-value < 0.05 was statistically significant.

## Results

### 1. BGN is highly enriched in high-grade glioma and correlated with poor prognosis

Based on data from TCGA, BGN expression in LGG and GBM are significantly dysregulated than normal brain tissue respectively (Fig. 1A, B). In the meantime, BGN expression was positively related with WHO grade (Fig. 1C). Prognostic value of BGN was also analyzed via log Rank test and showed that BGN high expressed group has poor prognosis than low expressed group with mean Hazard ratio (HR) 5.77, P<0.001(Fig. 1D). From Table. 1, we got the histopathological characteristics difference between.

**Table 1.**
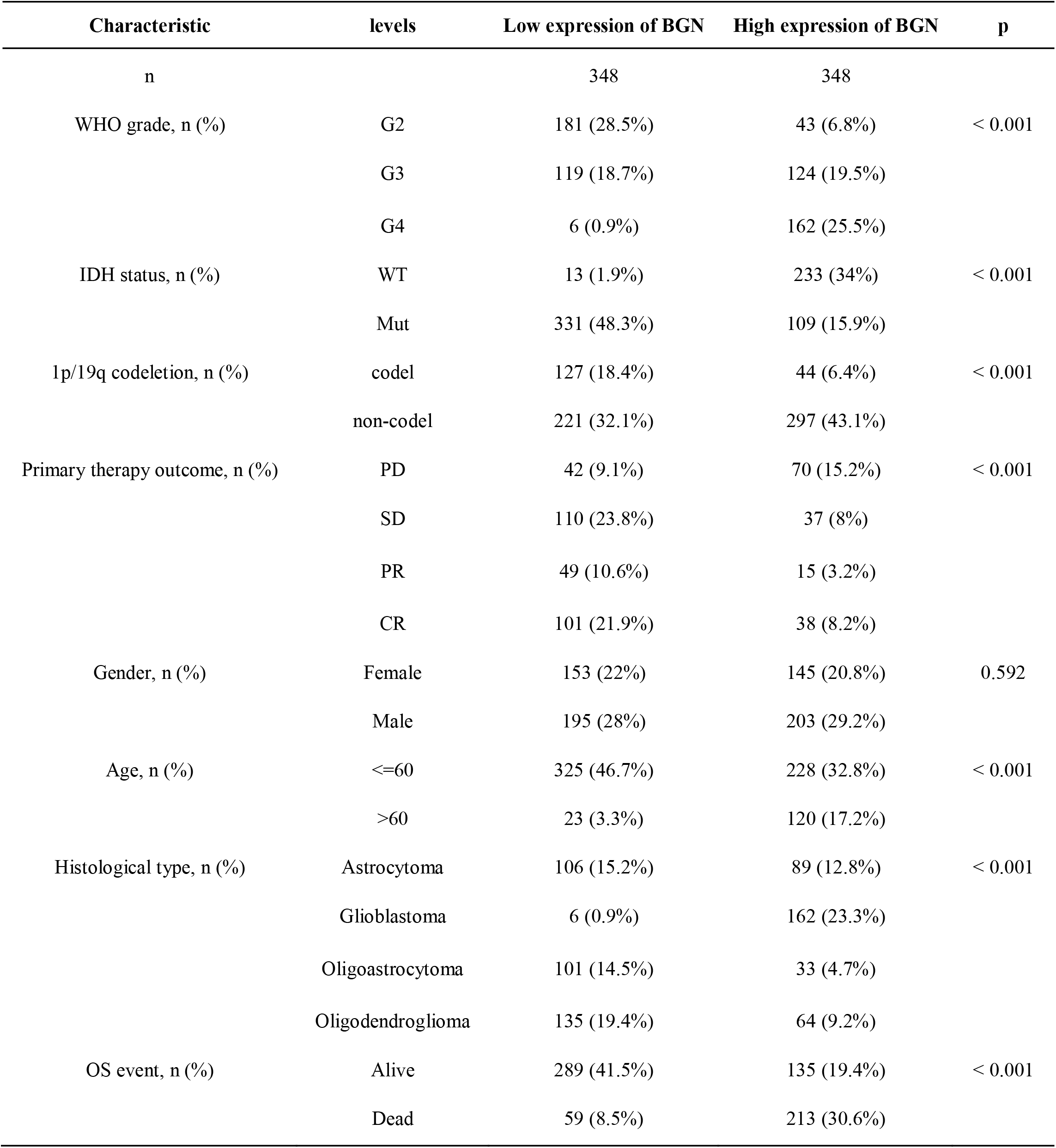
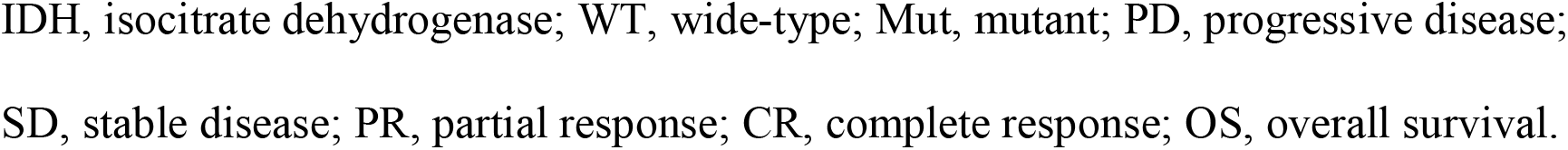
Relationship between BGN expression and clinicopathological characteristics in patients with glioma.

**Fig. 1.**
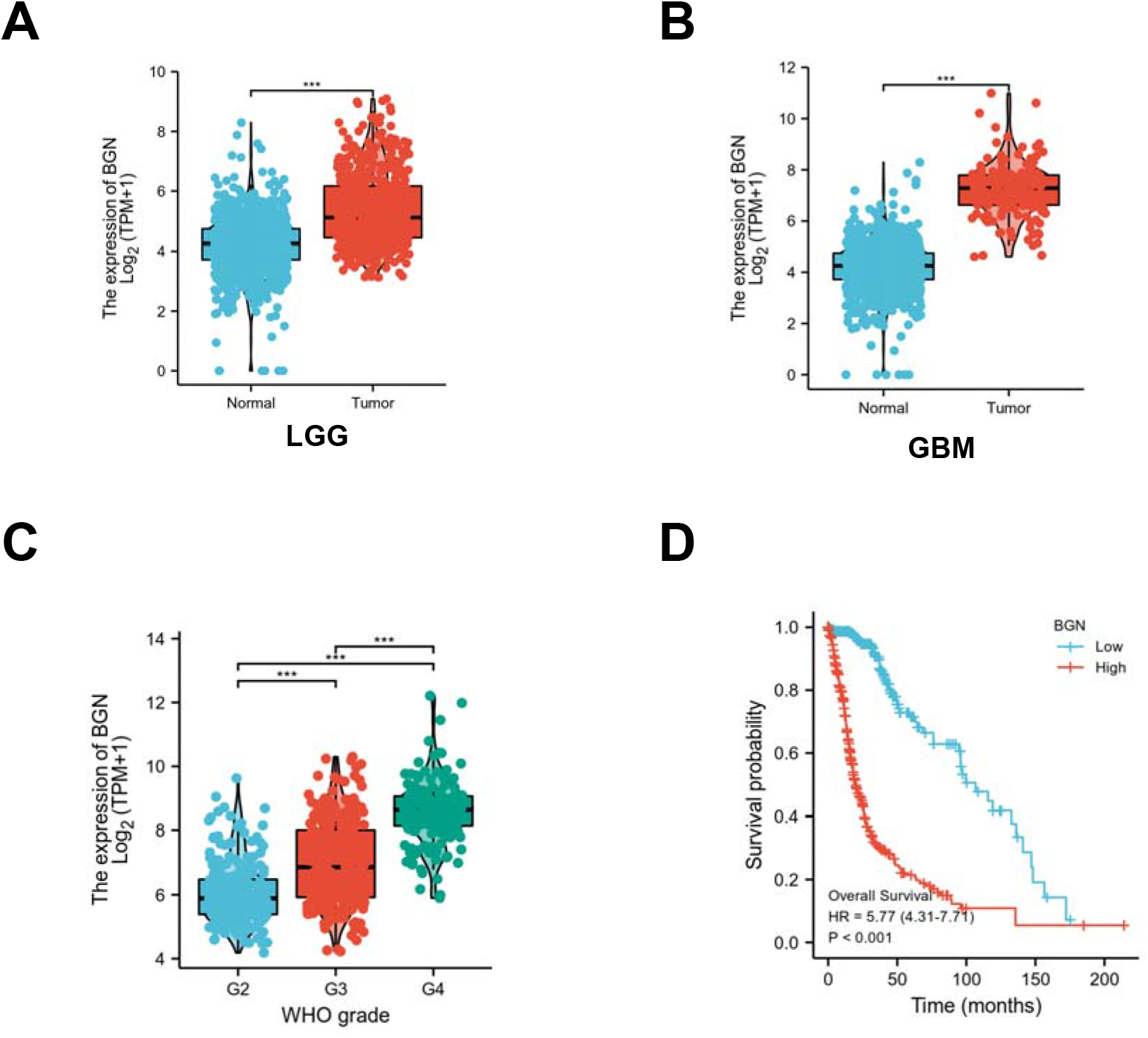
Expression of BGN in glioma. **(A-B)** The expression level of BGN in LGG and GBM and based on TCGA. **(C)** The expression level of BGN in different WHO glioma tissues. **(D)** Prognostic analysis in high and low BGN expression group

### 2. BGN promotes glioma cells EMT and invasion

Because of the relationship between EMT and TMZ drug resistance, the analysis of BGN and TMZ drug resistance was carried out.To investigate the function BGN in glioma progression, RNA sequencing data from TCGA were gone through R. Go and KEGG functional analysis showed BGN has an important role in ECM (Fig. 2A). Correlation between BGN and EMT-related gene expression in LGG and GBM showed that BGN might have relationship with cells EMT function (Fig. 2B). U251MG and HS683 cell line expressing different BGN level were establish by three siRNA (siRNA-a, siRNA-b, siRNA-c) and silenced glioma cells tended to express lower N-cadherin and VIM level (Fig. 2C), which means BGN participates in glioma EMT process. siBGN functional analysis showed a positive correlation with extracellular matrix construction and EMT, and Western blot and invasion experiments confirmed the results. Invasion assay was carried out to check glioma cells’ invasive ability under different BGN expression (Fig. 2D). Each group was repeat 5 times (Fig. 2E).

**Fig. 2.**
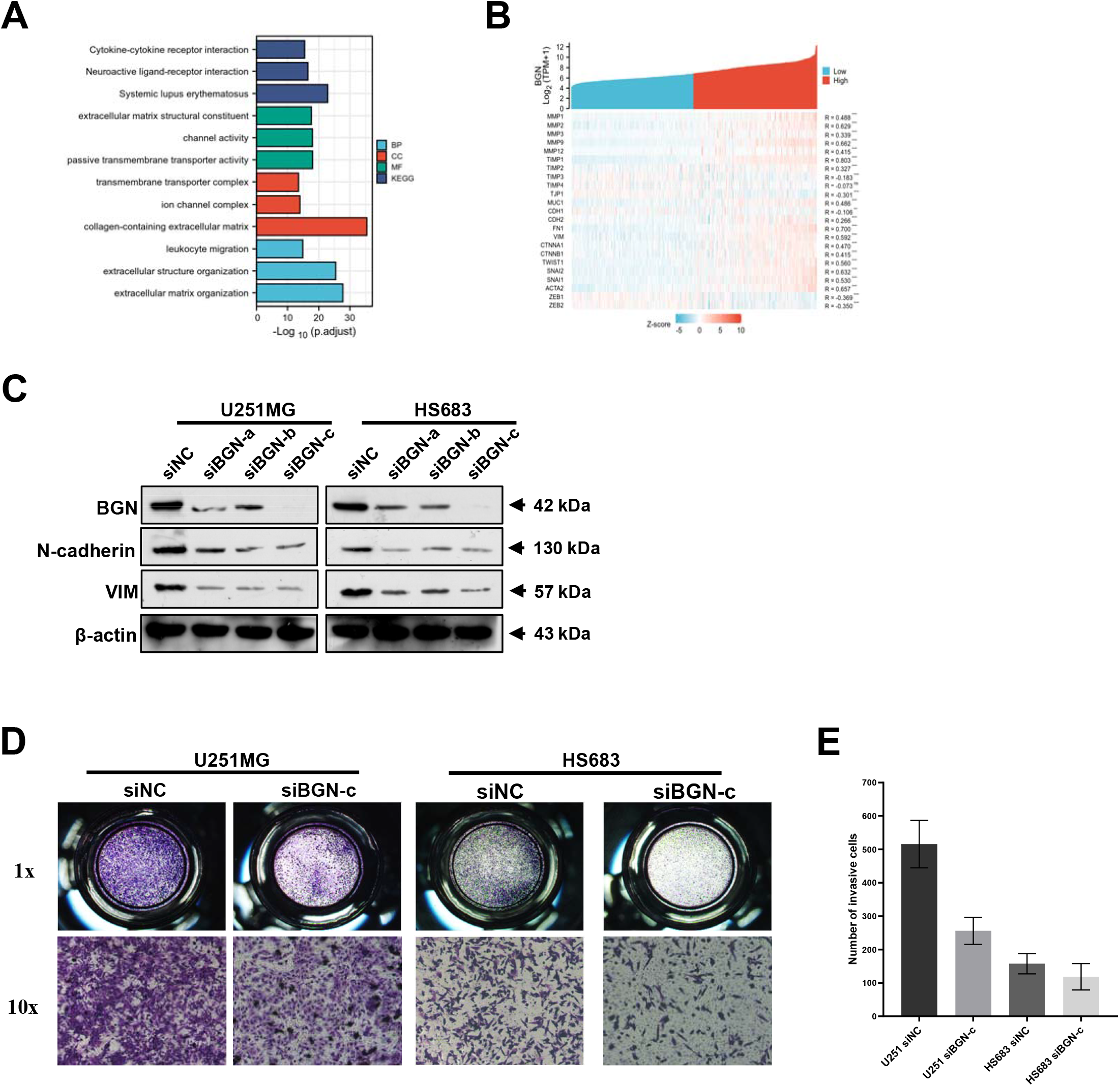
Enrichment results of GO and KEGG. **(A)** Go functional analysis BGN based on TCGA sequencing data. **(B)** Correlation between BGN and EMT-associated gene TCGA sequencing data. **(C)** Western blot results for BGN, N-cadherin and Vim expression after BGN silencing in U251MG and HS683 cells. **(D-E)** Invasive results in transwell chambers captured using Carl Zeiss Microscopy with both 1x and 10x objective magnification.

### 3. Silence of BGN sensitizes glioma to temozolomide

In light of the observed correlation between EMT and TMZ resistance in glioma cells [10], we conducted a Western blot analysis to assess BGN expression following treatment with 200µM TMZ for 4 hours, followed by 2 days of incubation in fresh complete medium. Remarkable upregulation of BGN expression was evident in both U251MG and HS683 glioma cells upon TMZ treatment compared to the mock-treated control group (Figure 3A). This intriguing finding prompted us to hypothesize that BGN may play a critical role in conferring TMZ resistance. To further explore this hypothesis, we conducted a TMZ treatment-based colony formation assay. Glioma cells were treated with 200µM TMZ and seeded at a density of 800 cells per 10cm dish. After 10 days of incubation to promote colony formation, the colonies were washed with PBS, fixated using 4% paraformaldehyde, and subsequently stained with violet crystal for visualization. It showed siBGN-a, siBGN-b, siBGN-c has less clonies than siNC group (Fig. 3B, C). Stemness marker expression after treated 200 µM for 4 hours. After TMZ treatment, siNC exhibit more expression of SOX2 and β-catenin proteins compared with siBGN-c group in western blot result (Fig. 3D).

**Fig. 3.**
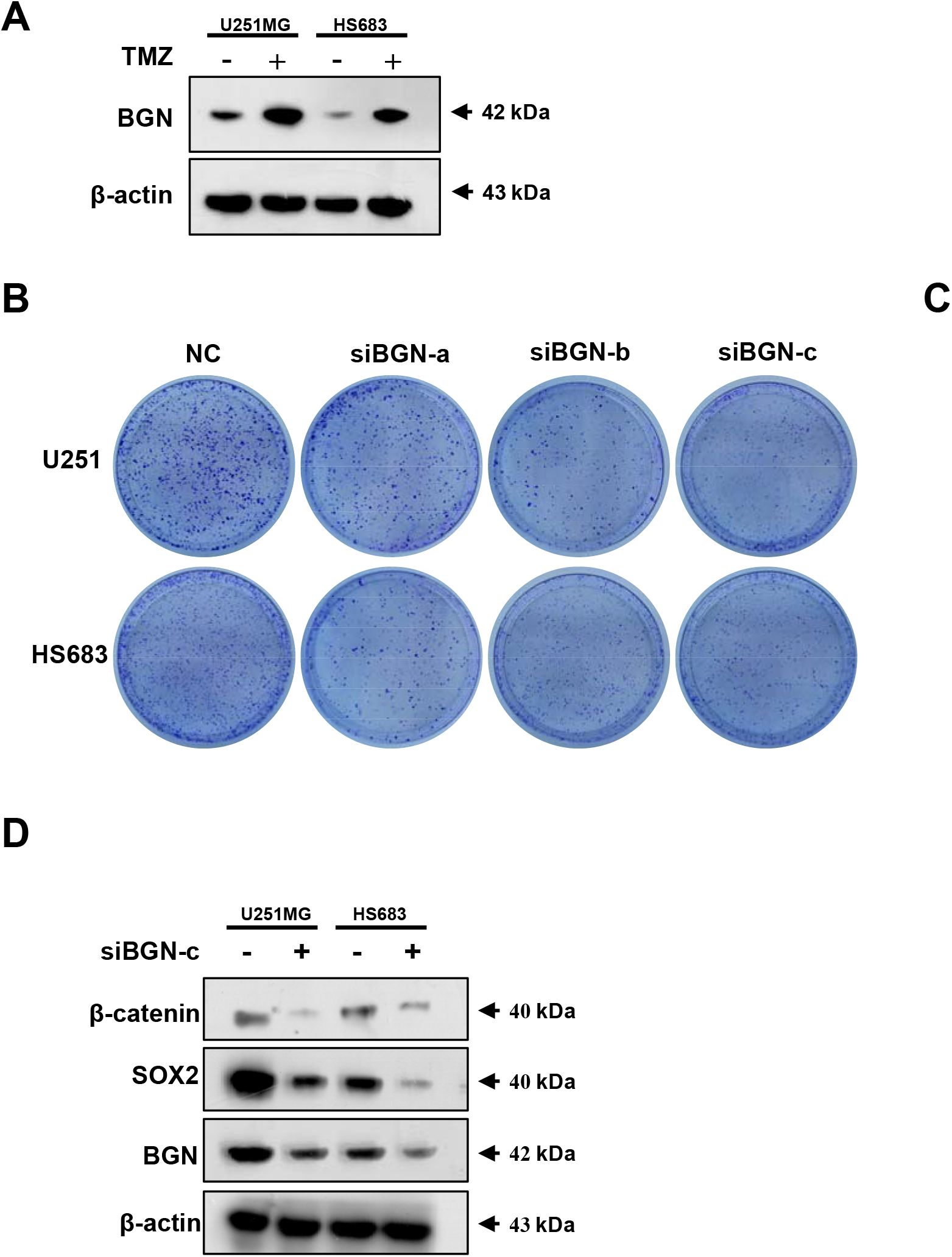
BGN expression and TMZ treatment. (A) BGN expression following treatment with 200µM TMZ for 4 hours, followed by 2 days of incubation in fresh complete medium. (B-C) U251MG and HS683 cells with different BNG expression were treated with 200µM TMZ for 4 hours and then seeded at a density of 800 cells per 10cm dish. After 10 days of incubation to promote colony formation, the colonies were washed with PBS, fixated using 4% paraformaldehyde, and subsequently stained with violet crystal for visualization (D) U251MG and HS683 cells were treated 200µM TMZ for 4 hours, followed by 2 days of incubation in fresh complete medium and subjected to western blot to check stemness markers expression (β-catenin and SOX2)

## Discussion

Gliomas are aggressive brain tumors associated with high mortality and treatment resistance[12]. In this study, we investigated the role of BGN in glioma progression and its potential as a prognostic marker and therapeutic target. Our analysis of data from The TCGA revealed that BGN expression is significantly dysregulated in low-grade gliomas LGG and glioblastoma GBM compared to normal brain tissue, suggesting its potential as a diagnostic biomarker for gliomas. Furthermore, we observed a positive correlation between BGN expression and the WHO grade of gliomas, indicating that BGN levels could serve as a valuable indicator of tumor aggressiveness and progression. Importantly, our results demonstrated that high BGN expression is associated with poor prognosis, making it a potential prognostic marker for glioma patients. Functionally, we investigated the role of BGN in glioma progression and found that BGN plays a crucial role in ECM regulation and EMT(Fig.1), processes known to contribute to glioma invasion and drug resistance[13]. ECM remodeling is associated with increased tumor invasion and metastasis, and our findings suggest that BGN may play a pivotal role in these processes. Moreover, EMT has been linked to increased invasiveness and drug resistance in glioma cells[14]. Our correlation analysis between BGN and EMT-related genes in both LGG and GBM provides evidence of a potential relationship between BGN and EMT function. To validate this, we conducted experiments using U251MG and HS683 glioma cell lines with different BGN expression levels achieved through siRNA silencing. The results revealed that silencing BGN led to decreased expression of N-cadherin and VIM, two key EMT markers[15], indicating that BGN might play a significant role in glioma EMT processes (Fig. 2). This suggests that BGN may contribute to the invasive and aggressive behavior of glioma cells through EMT regulation, positioning it as a potential therapeutic target to inhibit EMT and reduce glioma invasiveness. Additionally, we investigated the relationship between BGN and TMZ resistance in glioma cells. TMZ is a commonly used chemotherapeutic agent for glioma, but its efficacy is limited due to the development of drug resistance[16]. Our Western blot analysis revealed a remarkable upregulation of BGN expression in U251MG and HS683 glioma cells upon TMZ treatment, implying a potential involvement of BGN in TMZ resistance mechanisms. To further explore this hypothesis, we conducted a TMZ treatment-based colony formation assay, where glioma cells were treated with TMZ at a specific concentration. Interestingly, the silencing of BGN using siRNA resulted in reduced colony formation compared to the control group, suggesting that BGN may play a critical role in conferring TMZ resistance in glioma cells. Additionally, we assessed the expression of stemness markers SOX2 and β-catenin following TMZ treatment and observed that siRNA-mediated BGN silencing reduced their expression, which is consistent with their association with stemness and drug resistance (Fig. 3). These findings propose that BGN may contribute to TMZ resistance in glioma cells and indicate its potential as a therapeutic target to enhance the efficacy of TMZ treatment.

In conclusion, the results of our study underscore the clinical significance of BGN and pave the way for further investigations into the underlying molecular mechanisms to exploit BGN as a potential therapeutic target for glioma management.

## Ethical Statement

This study was approved by the appropriate Research Ethics Committee at the Beijing Tiantan Hospital (approval no:…?).

